# Golden Orbweavers Ignore Biological Rules: Phylogenomic and Comparative Analyses Unravel a Complex Evolution of Sexual Size Dimorphism

**DOI:** 10.1101/368233

**Authors:** Matjaž Kuntner, Chris A. Hamilton, Cheng Ren-Chung, Matjaž Gregorič, Nik Lupše, Tjaša Lokovšek, Emily Moriarty Lemmon, Alan R. Lemmon, Ingi Agnarsson, Jonathan A. Coddington, Jason E. Bond

## Abstract

Instances of sexual size dimorphism (SSD) provide the context for rigorous tests of biological rules of size evolution, such as Cope’s Rule (phyletic size increase), Rensch’s Rule (allometric patterns of male and female size), as well as male and female body size optima. In certain spider groups, such as the golden orbweavers (Nephilidae), extreme female-biased SSD (eSSD, female:male body length ≥ 2) is the norm. Nephilid genera construct webs of exaggerated proportions which can be aerial, arboricolous, or intermediate (hybrid). First, we established the backbone phylogeny of Nephilidae using 367 Anchored Hybrid Enrichment (AHE) markers, then combined these data with classical markers for a reference species-level phylogeny. Second, we used the phylogeny to test Cope and Rensch’s Rules, sex specific size optima, and the coevolution of web size, type, and features with female and male body size and their ratio, SSD. Male, but not female, size increases significantly over time, and refutes Cope’s Rule. Allometric analyses reject the converse, Rensch’s Rule. Male and female body sizes are uncorrelated. Female size evolution is random, but males evolve towards an optimum size (3.2–4.9 mm). Overall, female body size correlates positively with absolute web size. However, intermediate sized females build the largest webs (of the hybrid type), giant female *Nephila* and *Trichonephila* build smaller webs (of the aerial type), and the smallest females build the smallest webs (of the arboricolous type). We propose taxonomic changes based on the criteria of clade age, monophyly and exclusivity, classification information content, diagnosability, and arachnological community practice. We resurrect the family Nephilidae Simon 1894 that contains *Clitaetra* Simon 1889, the Cretaceous *Geratonephila* Poinar & Buckley 2012, *Herennia* Thorell 1877, *Indoetra* Kuntner 2006, new rank, *Nephila* Leach 1815, *Nephilengys* L. Koch 1872, *Nephilingis* Kuntner 2013, and *Trichonephila* Dahl 1911, new rank. We propose the new clade Orbipurae to contain Araneidae Clerck 1757, Phonognathidae Simon 1894, new rank, and Nephilidae. Nephilid female gigantism is a phylogenetically-ancient phenotype (over 100 ma), as is eSSD, though their magnitudes vary by lineage and, to some extent, biogeographically.

---

Evolution of body size is often attributed to biological laws. Rensch postulated phyletic increase in body size, i.e. size increase in evolutionary time, and referred to it as Cope’s Rule (Rensch 1948), now attributed to Depéret (Bokma et al. 2016). Authors disagree on whether the rule is generally valid or even biologically meaningful (Stanley 1973; Gould 1997), at what phylogenetic scale it may be applied (Heim and Knope 2015), and, importantly, how to test it (Hone and Benton 2005). Interpretations of this rule range from overall short-term fitness advantages of larger body size (Waller and Svensson 2017) to long-term size increases over geologic time (Hunt and Roy 2006).

Sexual size dimorphism (SSD) often seems to be correlated with extreme morphological, behavioral, and life history phenotypes in either sex. In female-biased size dimorphic organisms, SSD is defined as female-to-male body size ratio (or correlated body parts; Fairbairn 2007). Extreme, female biased values (≥ 2.0, termed eSSD) are rare in any animal group and eSSD provides a heuristic definition to identify extreme phenotypes (Scharff and Coddington 1997; Kuntner and Cheng 2016). In certain spider groups, eSSD is the norm, and values exceeding 5.0 are common (Hormiga et al. 2000). Such clades are the most extreme (eSSD) examples among all terrestrial animals (Kuntner and Elgar 2014), and thus figure prominently in studies of gendered body size evolution (Vollrath and Parker 1992; Coddington et al. 1997; Foellmer and Moya-Laraño 2007).

Prior studies have utilized nephilid and argiopine spider phylogenies to investigate patterns of sex-specific size evolution (Kuntner and Coddington 2009; Cheng and Kuntner 2014; Kuntner and Elgar 2014), but shared macroevolutionary patterns are scarce (Kuntner and Cheng 2016). For example, nephilid females and males were thought to grow larger phyletically, with the slope of female size evolution being steeper, thus eSSD was maintained (Kuntner and Elgar 2014). In argiopines (Araneidae), phyletic size change showed no net trend in either sex, and eSSD declined over time (Cheng and Kuntner 2014). Nephilids, but not argiopines, seem to follow Cope’s Rule. These prior studies of reconstructed size evolution concluded that argiopine size, nephilid female size, and SSD drifted randomly in time (i.e, they appear to follow a model of Brownian motion). Nephilid male size, however, fit a single optimum, under an Ornstein-Uhlenbeck model (Kuntner and Cheng 2016). Male and female nephilid sizes evolved independently (Kuntner and Coddington 2009; Higgins et al. 2011; Kuntner and Elgar 2014), but in argiopines male and female size were significantly correlated (Cheng and Kuntner 2014; Kuntner and Cheng 2016).

Rensch’s Rule predicts positive allometry for male versus female size in male-size dimorphic animals, either within species or clades, whereas the converse Rensch’s Rule predicts negative allometry in female-size dimorphic lineages (Abouheif and Fairbairn 1997; Fairbairn 1997; Blanckenhorn et al. 2007a). However, Rensch’s Rule has not been found for spiders at any phylogenetic level or fauna (Foellmer and Moya-Laraño 2007; Cheng and Kuntner 2014). Specifically, argiopines were isometric and nephilid sizes were uncorrelated (Kuntner and Cheng 2016).

As the ratio of gendered sizes, SSD is probably not a single trait under selection. SSD therefore is best considered as an epiphenomenon of potentially complex, taxon-specific, evolutionary changes in the size of each gender. The ratio is plausibly selected only if the direct interaction of males and females of different sizes affects fitness (Ramos et al. 2005; Lupše et al. 2016). Spider size variation in each gender can be caused by multiple proximate causes (Kuntner and Elgar 2014; Kuntner and Cheng 2016). Selection for larger, more fecund females, larger and stronger males, or for smaller, more agile males are all credible drivers of size evolution (Elgar 1991; Vollrath and Parker 1992; Head 1995; Higgins 2002; Moya-Laraño et al. 2002, 2009; Foellmer and Moya-Laraño 2007; Danielson-François et al. 2012; Cheng and Kuntner 2015). However, each factor may select for a different size, so that net selection on size could be equivocal (Kuntner and Elgar 2014).

Spider webs are, in an ecological sense, extensions of a spider’s phenotype and have evolved in diverse ways (Blackledge et al. 2011). Web characteristics can plausibly affect somatic trait evolution, but studies that statistically test links between body size and web characteristics across diverse species are rare (reviewed in Eberhard (1990)). Nephilids are ideal models for addressing such evolutionary questions (Table 1). The golden orb weavers, genus *Nephila*, are conspicuous tropical spiders (Kuntner 2017). Massive, colorful females construct their characteristic orb web with a golden shine, and these webs are unusually large (Kuntner et al. 2008). Tiny males, over 10 times smaller, and 100 times lighter than corresponding conspecific females (Kuntner et al. 2012), are less striking (Fig. 1a), and, notoriously, are often cannibalized by females (Elgar 1991; Schneider and Elgar 2001). *Nephila* and related genera (Kuntner et al. 2013) are popular lab animals and their biology is well understood (Fig. 1b-c). Because of female body size and web gigantism, nephilids have become models to study extreme phenotypes, especially their traits such as tough silk, large webs, eSSD, and sexually conflicted behaviors (Kuntner et al. 2009, 2016; Blackledge et al. 2011; Kuntner and Elgar 2014). Additionally, *Trichonephila clavipes* (formerly *Nephila;*Table 1) is the first orb web spider with an annotated genome (Babb et al. 2017). By revealing an unprecedented diversity of silk genes and their complex expression, this study suggested new directions of genetic and biomaterial research.

**Table 1.**
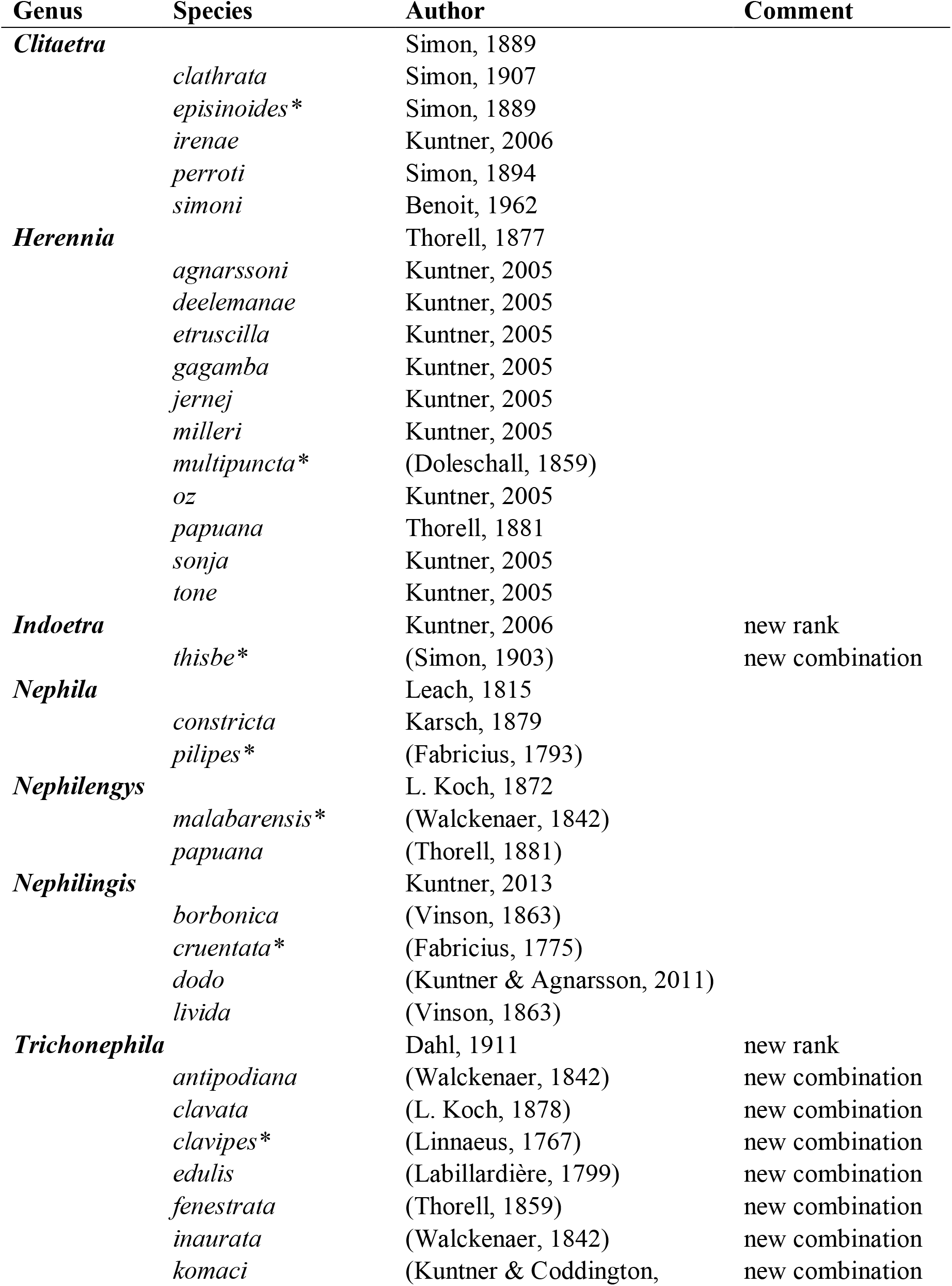
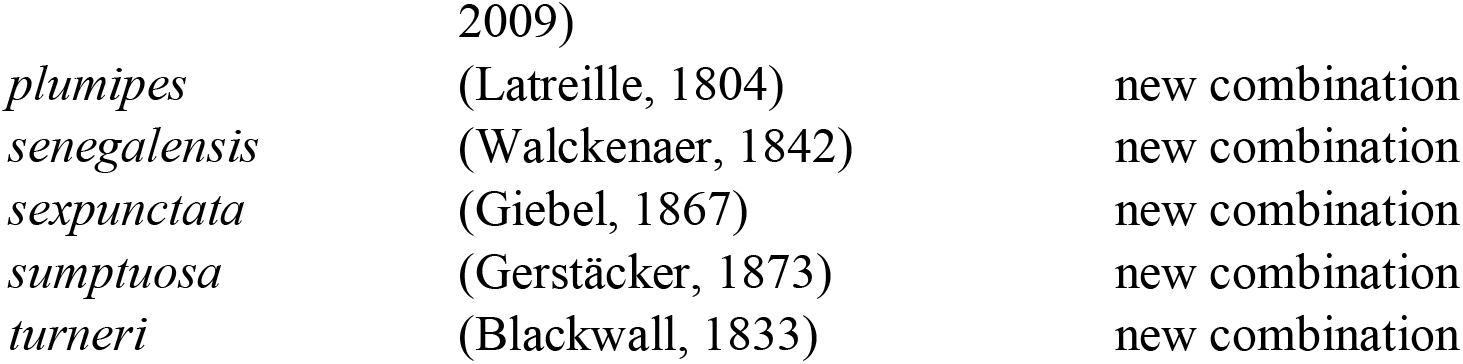
Currently valid contemporary genera and species of the family Nephilidae, including taxonomic changes proposed here. The genus *Nephila* includes additional names that are thought to be synonyms of *Nephila* or *Trichonephila* species listed below. Type species indicated by asterisks.

| Genus | Species | Author | Comment |
| --- | --- | --- | --- |
| ***Clitaetra*** |  | Simon, 1889 |  |
|  | *clathrata* | Simon, 1907 |  |
|  | *episinoides* * | Simon, 1889 |  |
|  | *irenae* | Kuntner, 2006 |  |
|  | *perroti* | Simon, 1894 |  |
|  | *simoni* | Benoit, 1962 |  |
| ***Herennia*** |  | Thorell, 1877 |  |
|  | *agnarssoni* | Kuntner, 2005 |  |
|  | *deelemanae* | Kuntner, 2005 |  |
|  | *etruscilla* | Kuntner, 2005 |  |
|  | *gagamba* | Kuntner, 2005 |  |
|  | *jernej* | Kuntner, 2005 |  |
|  | *milleri* | Kuntner, 2005 |  |
|  | *multipuncta* * | (Doleschall, 1859) |  |
|  | *oz* | Kuntner, 2005 |  |
|  | *papuana* | Thorell, 1881 |  |
|  | *sonja* | Kuntner, 2005 |  |
|  | *tone* | Kuntner, 2005 |  |
| ***Indoetra*** |  | Kuntner, 2006 | new rank |
|  | *thisbe* * | (Simon, 1903) | new combination |
| ***Nephila*** |  | Leach, 1815 |  |
|  | *constricta* | Karsch, 1879 |  |
|  | *pilipes* * | (Fabricius, 1793) |  |
| ***Nephilengys*** |  | L. Koch, 1872 |  |
|  | *malabarensis* * | (Walckenaer, 1842) |  |
|  | *papuana* | (Thorell, 1881) |  |
| ***Nephilingis*** |  | Kuntner, 2013 |  |
|  | *borbonica* | (Vinson, 1863) |  |
|  | *cruentata* * | (Fabricius, 1775) |  |
|  | *dodo* | (Kuntner & Agnarsson, 2011) |  |
|  | *livida* | (Vinson, 1863) |  |
| ***Trichonephila*** |  | Dahl, 1911 | new rank |
|  | *antipodiana* | (Walckenaer, 1842) | new combination |
|  | *clavata* | (L. Koch, 1878) | new combination |
|  | *clavipes* * | (Linnaeus, 1767) | new combination |
|  | *edulis* | (Labillardière, 1799) | new combination |
|  | *fenestrata* | (Thorell, 1859) | new combination |
|  | *inaurata* | (Walckenaer, 1842) | new combination |
|  | *komaci* | (Kuntner & Coddington, | new combination |

|  |  |  |
| --- | --- | --- |
|  | 2009) |  |
| *plumipes* | (Latreille, 1804) | new combination |
| *senegalensis* | (Walckenaer, 1842) | new combination |
| *sexpunctata* | (Giebel, 1867) | new combination |
| *sumptuosa* | (Gerstäcker, 1873) | new combination |
| *turneri* | (Blackwall, 1833) | new combination |

**Figure 1.**
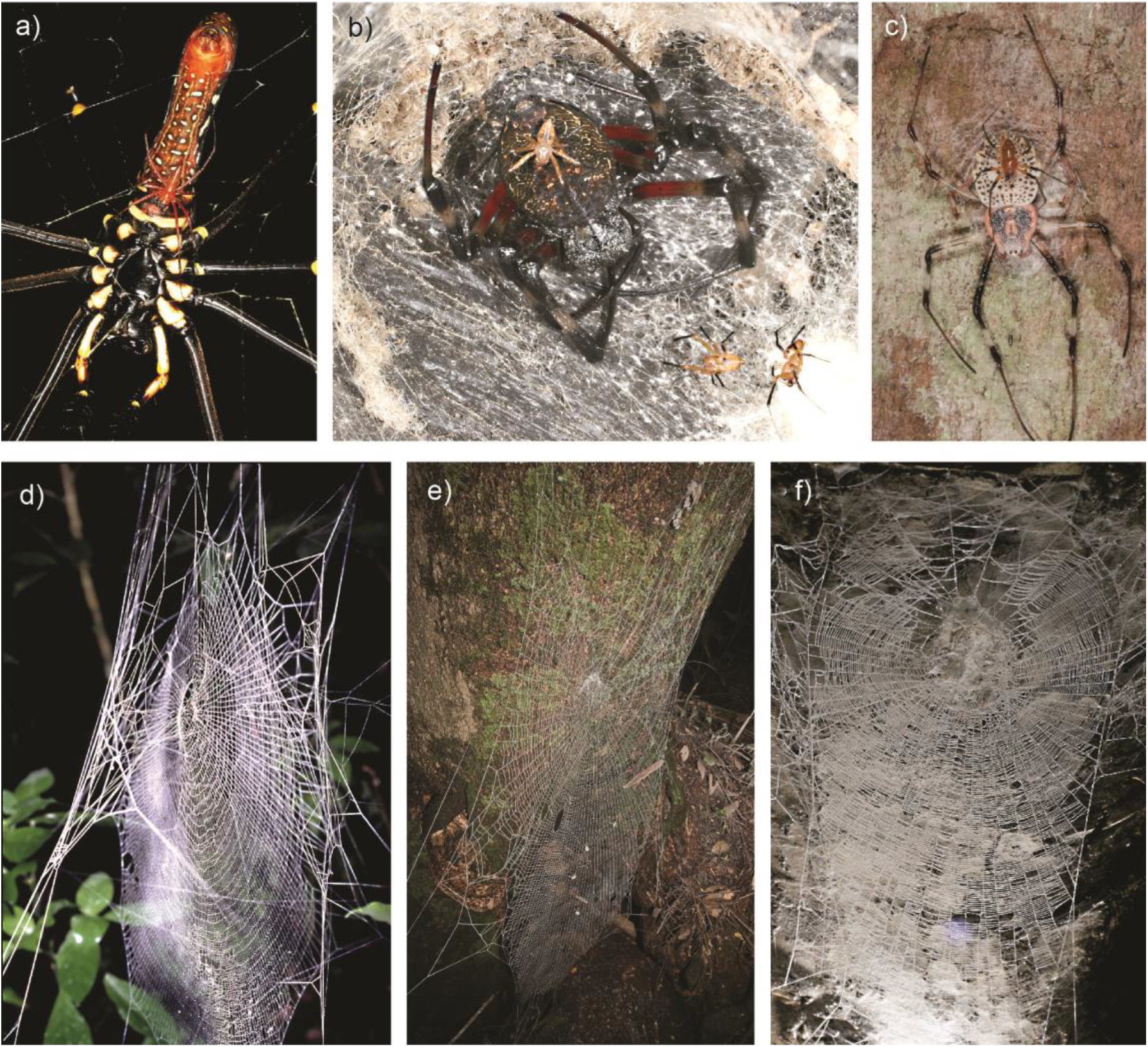
Representatives of nephilid genera, their sexual size dimorphism, and web types. (a) *Nephila pilipes* male on a female; (b) *Nephilingis cruentata* males accumulating around a female; (c) *Herennia multipuncta* male sitting on a female; (d) aerial orb web of *Trichonephila clavipes*; (e) hybrid web of *Nephilingis livida*; (f) arboricolous ladder web of *Clitaetra episinoides*.

The evolution of these remarkable traits continues to puzzle biologists, but well-formulated hypotheses are short-lived if phylogenetic estimates continue to change. A robust and resolved species-level phylogeny is crucial to understanding the evolution of extreme phenotypes, and to settle nephilid taxonomy and classification. Both published species-level phylogenies are outdated and potentially incorrect: one was based on morphological and behavioral phylogenetic data (Kuntner et al. 2008), and the other (Kuntner et al. 2013) on a few, commonly used mitochondrial and nuclear markers that, in the phylogenomics era, have clear limitations (Agnarsson et al. 2013).

The use of molecular data in spider phylogenetics has a relatively short history, with the first large studies starting to appear at the turn of the millennium (reviewed in Agnarsson et al. 2013). Until recently, research was necessarily limited to the few markers that could be amplified across spiders (Agnarsson et al. 2013; Dimitrov et al. 2017; Wheeler et al. 2017), which typically did a poor job recovering older nodes. The phylogenomics era brought forth transcriptomics and targeted sequence capture to begin more rigorously addressing many questions regarding spider evolution (Bond et al. 2014; Fernández et al. 2014, 2018; Garrison et al. 2016; Hamilton et al. 2016b; Starrett et al. 2017). Many formerly stable hypotheses have not survived this data revolution, including the monophyly of orb-weavers, relationships of primitive araneomorphs, patterns of spider diversification, and ages of major spider groups (Bond et al. 2014; Fernández et al. 2014; Garrison et al. 2016; Maddison et al. 2017). While relationships among spider families seem to be stabilizing, few species-level phylogenomic studies have been published (Hamilton et al. 2016a). Such phylogenies are necessary to test detailed comparative hypotheses.

Here we use Anchored Hybrid Enrichment (AHE) methodology (Lemmon et al. 2012) to provide a well corroborated species-level phylogeny and to estimate lineage ages. We test the reciprocal monophyly of Nephilidae *sensu* Kuntner (2006) and Kuntner et al. (2013) and its genera using 22 ingroup taxa, then use this constrained backbone phylogeny to place an additional nine nephilid species using the classical markers (Kuntner et al. 2013) (total nephilid diversity ∼37 spp.). We use this nephilid topology to test hypotheses on body and web form and size evolution, detailed below. We also infer nephilid age using the 97–100 Ma monotypic Cretaceous Burmese amber *Geratonephila burmanica* Poinar 2012 as a constraint. The phylogeny represents the foundation to correct and refine the taxonomy of Nephilidae (Table 1). And finally, given the new phylogeny, we pose three broad questions: 1) Do male and female size evolve independently, do they evolve towards optima, and how does their evolution affect SSD?; 2) Do nephilids obey Cope’s Rule and the converse Rensch’s Rule?; 3) What is the relationship among spider body size and web size, web types, web architecture, and SSD?

## Materials & Methods

### Phylogenomics

We employed the AHE targeted-sequencing approach for spiders (outlined in Hamilton et al. 2016b) to target 585 single copy orthologous loci from across the genome. These loci have been shown to possess sufficient variation for resolving both shallow and deep-scale evolutionary relationships throughout the Araneae. Hamilton et al. (2016b), Maddison et al. (2017), and Godwin et al. (2018) have used AHE to recover genus and species-level relationships within spider families, Theraphosidae, Salticidae, and Halonoproctidae/Ctenizidae.

We obtained sequence data for 22 nephilids and 11 outgroups. High-quality genomic DNA (≥1µg) for all specimens was extracted from leg tissue stored in ≥95% EtOH at –80° C, using an optimized protocol on the MagMAX Express magnetic particle processor robot (Vidergar et al. 2014). DNA concentration was evaluated through agarose gel electrophoresis and spectrophotometry using a NanoDrop ND-1000.

AHE data, including library preparation, enrichment, and sequencing, were generated at the Center for Anchored Phylogenomics at Florida State University (http://www.anchoredphylogeny.com) following Lemmon et al. (2012), Prum et al. (2015), and Hamilton et al. (2016b). Up to 500 ng of each DNA sample was sonicated to a fragment size of ∼300–800 bp using a Covaris E220 ultrasonicator. Indexed libraries were then prepared following Meyer and Kircher (2010), but with modifications for automation on a Beckman-Coulter Biomek FXp liquid-handling robot (see Hamilton et al. (2016b) for details). Size-selection was performed after blunt-end repair using SPRI select beads (Beckman-Coulter Inc.; 0.9x ratio of bead to sample volume). Indexed samples were pooled at equal quantities (16 samples per pool), and then each pool was enriched using the AHE Spider Probe kit v1 (Hamilton et al. 2016b) and a modified v2 (Hamilton et al, unpublished), which refines the previous v1 capture probes to capture the same loci but yield greater enrichment within araneomorph spiders. After enrichment, the two reactions were pooled in equal quantities and sequenced on one PE150 Illumina HiSeq 2500 lane (35.2Gb) at Florida State University Translational Science Laboratory in the College of Medicine. Prior to assembly, overlapping paired reads were merged following Rokyta et al. (2012). For each read pair, the probability of obtaining the observed number of matches by chance was evaluated for each possible degree of overlap. The overlap with the lowest probability was chosen if the p-value was less than 10^-10^, a stringent threshold that helps avoid chance matches in repetitive regions (Rokyta et al. 2012). Read pairs failing to merge were utilized but left unmerged during the assembly. Subsequent bioinformatic pipelines (data processing, sequence assembly, quality control, orthology search, alignment) follow Hamilton et al. (2016b), with contigs derived from fewer than 20 reads being removed before orthology assessment. Alignments were performed in MAFFT v. 7 (Katoh and Standley 2013) with gaps treated as missing characters.

We defined two AHE datasets, based on “strict” and “loose” trimming/masking thresholds (“strict”: goodSites=14, propSame=0.5, missingAllowed=5; “loose”: goodSites=16, propSame=0.6, missingAllowed=11; see Hamilton et al. (2016b) for details) in order to evaluate matrix occupancy on phylogeny estimation. The “strict” dataset had 206 loci (42,396 bp total sites; 13,338 informative sites) with 9% missing data, while the “loose” had 367 loci (89,212 bp total sites; 27,129 informative sites) with 22% missing data, each with the same 34 taxa. We partitioned the data by locus and concatenated, with the resulting supermatrix analyzed using Maximum Likelihood in IQ-TREE v1.4.2 (Nguyen et al. 2015) using the–m TEST command and 1000 rapid bootstraps. IQ-TREE analyses allow for a much larger suite of evolutionary models to be tested for and applied per locus/partition. RogueNaRok (Aberer et al. 2013) was used to investigate the presence of rogue taxa and whether they were influencing any part of the inference or topology. In addition to analyzing the supermatrix data, we employed ASTRAL v4.10.12 (Mirarab and Warnow 2015), a genome-scale coalescent-based species tree estimation, on both “loose” and “strict” individual gene trees inferred using IQ-TREE, and employing the same parameters as the supermatrix (above). We evaluated the AHE topologies for topological congruence/disagreement and clade support, selecting a representative AHE topology to be used as the backbone constraint. In order to provide additional ingroup taxa (Kuntner et al. 2013) for which AHE data were not available, a chimaeric dataset was created by merging the “strict” dataset with a legacy dataset of three standard loci from previous Nephilidae phylogeny inference (*cox1*, 16S rRNA, ND1; Kuntner et al. 2013), where all tips included these 3 loci. A simplified backbone phylogeny was created (“strict” and “loose” inferences produced identical topologies), by stripping branch lengths and simplifying names. This tree was then used as a constraint for the chimaeric inference. The total dataset comprised 209 loci (44,579bp) for 45 taxa. The combined “constraint” analysis placed a total of 31 nephilid species within a reference, species-level phylogeny, and was inferred using IQ-TREE with the same parameters as above. All consensus sequences, alignments, tree files, and scripts are available on the Dryad Data Repository (doi: to be added).

### Divergence Estimation

Lineage divergence times were estimated using the RelTime maximum likelihood method explicitly proposed for dating nodes from large phylogenomic datasets (Tamura et al. 2012). In order to evaluate consistency, divergence times were estimated for both the 367 loci (“loose”) supermatrix, to include more putatively informative sites for dating, and the 206 loci (“strict”) dataset. The “constraint” dataset was not evaluated due to increases in missing data for those taxa that were added. Each ML best tree was used as the reference topology for each respective analysis; topologies were identical between “strict” and “loose”. Local clocks were used for each lineage, with no clock rates merged. The HKY substitution model was employed with gamma distributed rates among sites (and 5 discrete gamma categories). Partial deletion was used, with a cutoff of 50% missing data at a site.

Due to the lack of available or informative fossils, we explored a suite of different program parameters and fossil calibration min/max boundaries (see Discussion for more detail). Two nodes were calibrated. To set a minimum age on the *Trichonephila* clade, a *Nephila* species in Dominican amber was used (Wunderlich 1986). Because we cannot assume when a lineage might have split or gone extinct, we set a hard minimum of 16 Ma (the age of the fossil), and a softer maximum age of 23 Ma on the *Trichonephila* clade – based on the close resemblance of a Dominican amber “*Nephila”* species to the contemporary *Trichonephila* (Wunderlich 1986). This boundary was an attempt to account for date flexibility, aging the fossil slightly older than the rock where it was recovered.

*Geratonephila burmanica* Poinar 2012, from early Cretaceous Burmese amber, is thought to be an ancestral nephilid (Poinar and Buckley 2012). The strongest nephilid synapomorphy is striae on the cheliceral boss (Kuntner et al. 2008), but this feature is not mentioned or visible in the *Geratonephila* description – cheliceral striae are rarely visible in any amber specimen. Striae are difficult to see even in extant male nephilids, and, in any case, our request to examine the type was ignored. The palpal morphology resembles a nephilid because the embolic conductor fully encloses the embolus, the cymbium is cup shaped, the paracymbium has an apophysis, and the bulb lacks the tegular apophyses typical of araneids and phonognathids. In fact, the *G. burmanica* palp closely resembles *Clitaetra episinoides* from the Comoros (compare Poinar and Buckley’s (2012) Figure 3 with Kuntner’s (2006) Figure 13). Both have globular cymbia mislabeled as the tegulum by Poinar and Buckley (P&B), similar paracymbia (not labeled by P&B), tegula (mislabeled as subtegulum by P&B), embolic bases (not labeled by P&B), and embolic conductors (not labeled by P&B). The *Geratonephila* tegulum and the embolic conductor seem to be rotated out of their usual position. *Geratonephila* could be an early offshoot of *Clitaetra*, but since we were unable to verify the type specimen, we conservatively treat it as a stem nephilid.

To set an age for the Cretaceous *Geratonephila*, a hard minimum boundary of 97 Ma (age of the Burmese amber) was thus set, as well as a softer maximum boundary at 146 Ma, the beginning of the Cretaceous. If *Geratonephila burmanica* was treated as a stem *Clitaetra* at 146–97 Ma, the dated splits would be vastly older than those in published phylogenomic analyses (Bond et al. 2014; Garrison et al. 2016). Our more conservative dating scheme placing *Geratonephila* at the stem of the Nephilidae seems to be better justified.

### Body Size

We measured the total body length, carapace length and width, and first leg patella + tibia length for a total of 480 males and females of the 28 nephilid species that are known from both sexes. We calculated SSD as the ratio of female to male values for the above measurements (Table S1).

### Web Types and Size

Nephilids spin three major types of webs. *Nephila* and *Trichonephila* spin large (often ≥ 100 cm diameter), completely aerial orb webs (Fig. 1d) (Kuntner et al. 2008; Kuntner 2017). The nephilid aerial web differs from typical orb architecture (e.g. Araneidae and Tetragnathidae) in details of radii, frames, and spirals, but especially in its asymmetrically placed hub. In contrast, the arboricolous ladder web of *Herennia* and *Clitaetra* (Fig. 1f) is least similar to orbs. These ladder-shaped webs are spun only on tree trunks. They have parallel (rather than converging) side frames attached to the trunk, relatively horizontal, parallel “spirals,” and hubs that attach to the substrate (Kuntner 2005, 2006). The third architecture, typical of *Nephilengys* and *Nephilingis*, is intermediate between the first two, neither fully aerial nor substrate bound (“hybrid”; Fig. 1e). The upper frames and hubs attach to substrate (e.g. trees or house roofs), but their aerial capture areas are rich in radii and spirals (Kuntner 2007; Kuntner and Agnarsson 2011; Kuntner et al. 2013). Hybrid webs vary in size but can be extremely large: *Nephilingis livida* webs are up to 151 cm high, easily surpassing *Nephila* or *Trichonephila* aerial webs (up to 116 cm), and *Herennia* ladders (up to 100 cm). The heavy female spiders hide in substrate retreats during the day.

In the field, we measured three web parameters: the (a) horizontal and (b) vertical diameters, as well as the distance from the hub to the top edge (c) (Kuntner et al. 2010) for 18 species (Table S2). We calculated web area (WA) using the formula WA=(a/2)*(b/2)*π (Blackledge and Gillespie 2002; Gregorič et al. 2011), the hub displacement index (HD) using the formula HD=(b-c)/b (Kuntner et al. 2008), and the ladder index (LI) using the formula LI=b/a (Peters 1937; Kuntner et al. 2008). Larger values of HD and LI imply more asymmetric webs.

### Comparative Analyses

Using the reference topology, pruned to include only the ingroup Nephilidae, we analyzed data on body size, web area and type in a comparative framework. We inferred ancestral body size and SSD in Mesquite v.3.04 (Maddison and Maddison 2015) under squared change parsimony. We used phylogenetically independent contrasts (PIC) in Mesquite to calculate correlations in continuous data. To test Cope’s rule, we used a linear regression model that regressed ancestral sizes against cladogenetic events. To test for Rensch’s Rule, we performed an allometric analysis (model II regression analysis) using the function “lmodel2” in the R package “lmodel2” (Legendre 2014) with the phylogenetically independent contrasts of log10-transformed body sizes. We ran major-axis regression of male body size on female body size with 10,000 simulations. To test optimum size evolution, we fitted three evolutionary models (Brownian motion versus single optimum Ornstein-Uhlenbeck versus Brownian motion with a directional trend) on nephilid size data using the function “fitContinuous” in the R package “geiger” (Harmon et al. 2008). We selected the best fit model using a likelihood ratio test. To examine the relationships between female web characteristics and the size of both sexes and web types, we employed Bayesian analyses of generalized linear mixed models, with phylogeny as a random factor, via the function “MCMCglmm” in the R package “MCMCglmm” (Hadfield 2010). We treated WA, HD and LI as dependent variables, and female size and male size, as well as web type, as independent variables. If regression revealed significant differences between web types, additional multiple comparisons of dependent variables among the three web types were calculated. Finally, we tested if body size and SSD differed among web types via “MCMCglmm” analyses. Appendix S1 provides the R code for all above analyses.

## Results

### Phylogeny

AHE phylogenomics inferred robust support for the relationships of 22 nephilid species (Fig. 2a). All analyses on the concatenated data (Fig. S1) (“strict” or “loose”), as well as species trees (Fig. S2), agree on nephilid monophyly, generally with robust bootstrap support throughout. Additionally, no taxa were discovered to be influencing the phylogenetic inference. *Nephilengys, Herennia*, and *Nephilingis* are monophyletic and confined to well defined biogeographical regions: *Nephilengys* + *Herennia* is Australasian, *Nephilingis* is Afrotropical. *Clitaetra* is also monophyletic (but only represented by two Afrotropical species). Formerly, species recognized as *Trichonephila* (Table 1) were recognized as *Nephila*, but Fig. 2a shows that this classical *Nephila* is diphyletic. These genera, as well as the topological position of the true *Nephila* (i.e, sister to all other Nephilidae), are fully supported in all analyses, the next distal clade separating *Clitaetra* from the remaining nephilids is strongly supported in the concatenated analyses (Fig. 2a; Fig. S1), but relatively weakly in ASTRAL analyses (Fig. S2).

**Figure 2.**
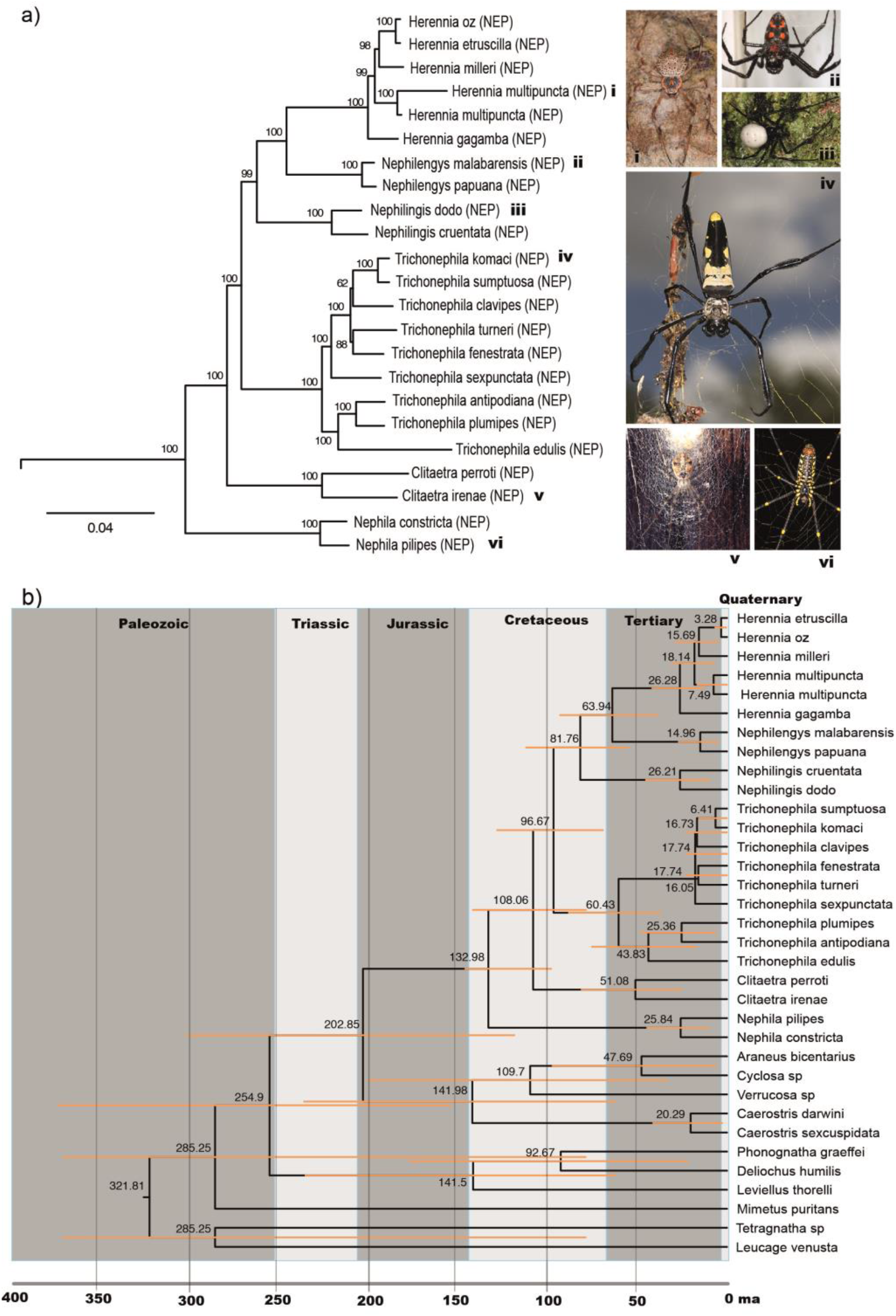
Anchored Hybrid Enrichment (AHE) resolves the relationships of 22 nephilid species. (a) Maximum Likelihood inference of concatenated data from 367 loci (“loose” dataset); This tree shows the ingroup, with field photographs of typical representatives; For entire and other topologies, see Figures S1, S2, S3. (b) Chronogram on the same taxon and AHE sample with labeled node ages in million years (ma) and their 95% confidence intervals. NEP = Nephilidae.

The AHE data (Figs. S1, S2, S3) supports the monophyly of three major araneoid clades, nephilids (NEP), araneids (ARA; represented here by *Araneus, Cyclosa, Verrucosa, Caerostris*), and phonognathids (PHO; represented by *Phonognatha, Deliochus, Leviellus*). These clades are consistently well supported, and, if combined, could be considered as Araneidae s.l. (Dimitrov et al. 2017). However, the sister relationships among the three vary, with most concatenated data recovering NEP + ARA (Fig. S1, S3) and most ASTRAL analyses recovering NEP + PHO (Fig. S2), but always with low support. No analyses support ARA + PHO.

Lastly, we added nine nephilid species that lacked AHE data, but for which three loci (*cox1*, 16S rRNA, ND1) were available (Kuntner et al. 2013). Using the AHE topology as a constraint, we placed *Herennia papuana* and *H. tone, Nephilingis borbonica* and *N. livida, Trichonephila inaurata, T*. *senegalensis*, and *T*. *clavata, Clitaetra episinoides* and *Indoetra thisbe*. The maximum likelihood analyses resulted in a robust and resolved tree, our reference species-level phylogeny (Fig. S4). Because the topology does not diverge from our AHE data, we converted this topology to an undated, relative rates ultrametric tree (Fig. 3) for comparative analyses.

**Figure 3.**
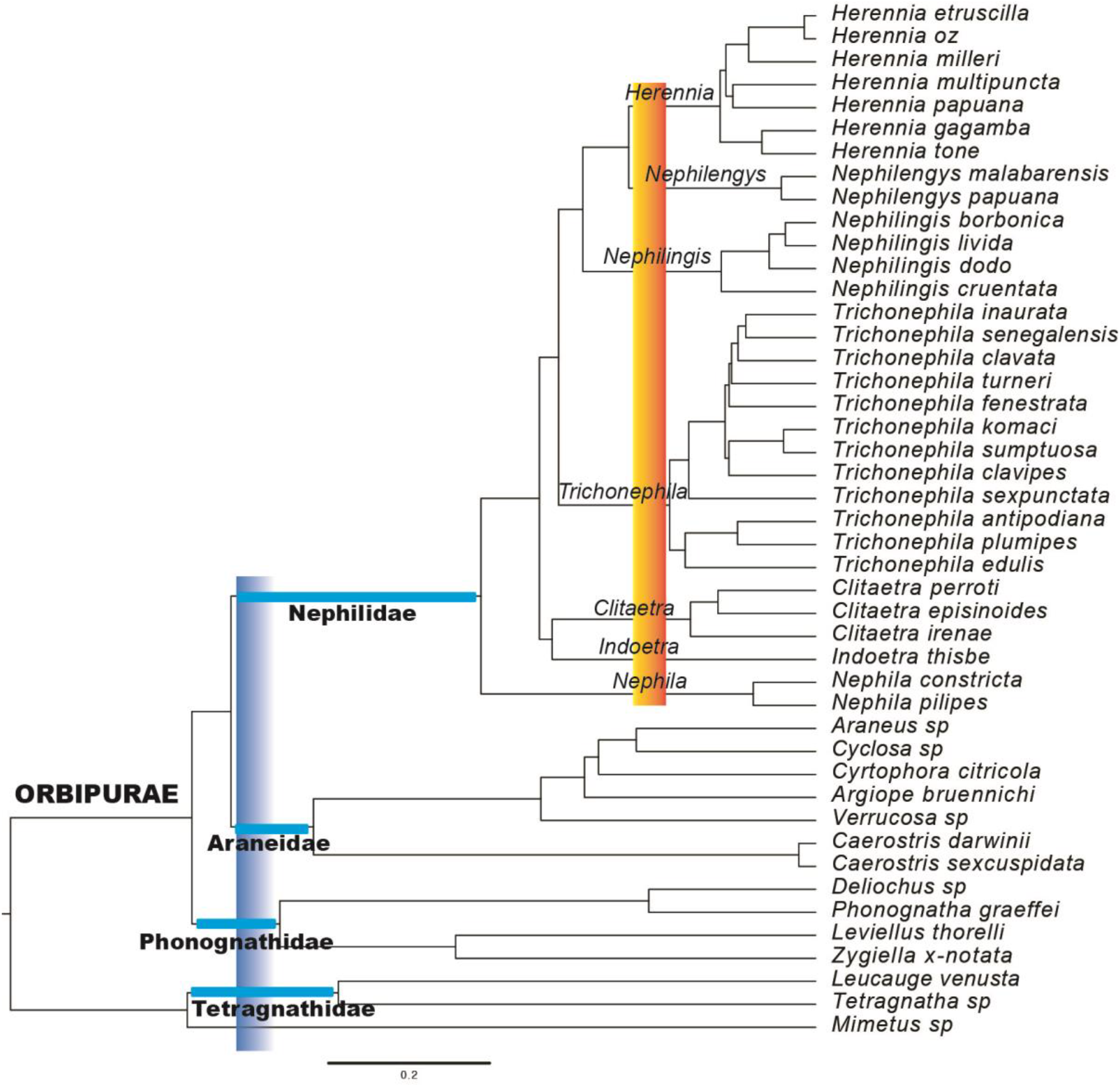
Phylogeny of Nephilidae with taxonomic implications. Ultrametricized tree is from a constrained, species-level phylogeny (see Fig. S4). The clade age taxonomic criterion aligns the families Nephilidae, Araneidae, Phonognathidae, and Tetragnathidae (blue vertical bar), as well as the seven nephilid genera (orange vertical bar).

### Fossil Evidence and Divergence Time Estimation

Time calibrated analysis using *Geratonephila* places the origin of nephilids well into the Cretaceous, estimated at 133 Ma (range 97 – 146) (Fig. 2b). *Herennia* is found to have originated around 26 Ma (13 – 42), *Nephilengys* around 15 Ma (5 – 27), *Nephilingis* around 26 Ma (10 – 45), *Trichonephila* around 60 Ma (36 – 88), *Clitaetra* around 51 Ma (25 – 81), and *Nephila* around 25 Ma (10 – 45). These estimated ages are older than prior analyses (Kuntner et al. 2013), a common trend in the age of phylogenomics. The origins of Araneidae and Phonognathidae are both estimated at around 140 Ma, although with large confidence intervals (Fig. 2b) and low taxonomic sampling.

### Taxonomy

Our primary rationale for taxonomic decisions are monophyly and estimated node age (Fig. 2b). Ages of all genera are comparable (Fig. 3, orange vertical bar). Table 1 summarizes the taxonomic changes listed below. Because the former, classic *Nephila* is diphyletic, (Fig. 2a), *Nephila* Leach, 1815 includes only its type, the Australasian *N. pilipes* and the African *N. constricta*. We assign the remaining 12 species to the circumtropical *Trichonephila* Dahl, 1911, new rank (formerly a *Nephila* subgenus; Dahl 1911). As its type species, we designate *Aranea clavipes* Linnaeus 1767. We elevate *Indoetra* Kuntner 2006, new rank, to genus (formerly a subgenus of *Clitaetra*; Kuntner 2006), and designate as its type species *I. thisbe* new combination from Sri Lanka. *Clitaetra* now includes the African and Western Indian Ocean island fauna.

For the same reasons (see also Discussion), we formally recognize Nephilidae as a family (Fig. 3, blue vertical bar). Kuntner (2006) elevated Nephilidae Simon 1894 to family rank and defined it phylogenetically as the least inclusive clade containing *Clitaetra, Herennia, Nephila* and *Nephilengys*. That definition also includes *Trichonephila, Nephilingis* and *Indoetra*. A striated cheliceral boss, an unreversed synapomorphy, diagnoses Nephilidae (Kuntner 2006). The family Araneidae Clerck 1757 is defined as the least inclusive clade containing *Araneus, Argiope, Caerostris, Cyclosa, Cyrtophora*, and *Verrucosa*. The family Phonognathidae Simon 1894 new rank is defined as the least inclusive clade containing *Deliochus, Leviellus, Phonognatha*, and *Zygiella*. Previously treated as Zygiellidae Wunderlich 2004, or as Zygiellinae (Hormiga and Griswold 2014; Gregorič et al. 2015; Dimitrov et al. 2017; Kallal and Hormiga 2018; Kallal et al. 2018), the family group name Phonognathidae (Simon 1894 as Phonognatheae) has precedence. Wunderlich (2004) based Zygiellidae on Zygielleae Simon 1929, with type genus *Zygiella* F.O.P. Cambridge 1902, when Zygiellidae did not contain *Phonognatha*. Because *Phonognatha* is now part of this clade, as are *Deliochus* and *Artifex* (Kallal and Hormiga 2018), the oldest family group name prevails (ICZN, Article 23.3).

If nephilids (and phonognathids) are included in Araneidae *s. l*. as proposed by works referenced above, that family becomes extraordinarily old and morphologically complex compared to other spider families (see Orbipurae, Fig. 3). If Nephilidae, Araneidae, and Phonognathidae are recognized as families (Fig. 3, blue horizontal and vertical lines), their ages, morphology, and phylogenetic distinctiveness are comparable to other spider families (Garrison et al. 2016). Additionally, Araneidae, Nephilidae, and Phonognathidae are reciprocally monophyletic. We propose the rankless name Orbipurae (from orb + pure or classic, a Latinized feminine plural) for this clade, defined as the least inclusive clade containing *Nephila pilipes* (Fabricius 1793)*, Araneus angulatus* Clerck 1757, and *Phonognatha graeffei* (Keyserling 1865).

### Body Size, SSD and Web Evolution

Table S1 provides average sizes of male and female body parts for 28 species (n = 480 specimens). The average nephilid total female body length is 20.4 mm (range 3.5 – 36.1) and male body length is 4.2 mm (2.5 – 7.1). The average SSD is 5.0 (1.4–11.4).

Table S2 summarizes female web features for 18 of the 28 species in Table S1. A parsimony reconstruction of web types (Fig. 4a) suggests that the aerial web is ancestral in nephilids, and thus homologous in *Nephila* and *Trichonephila*. *Herennia, Clitaetra* and *Indoetra* have arboricolous webs. *Clitaetra* and *Indoetra* webs are planar with space between the tree surface and the web, but *Herennia* webs are convex, following tree trunk curvature, with a hub cup attached to the tree. Interestingly, these two arboricolous web types evolved independently (Fig. 4a). Their natural history on tree trunks is similar, but convergent. Hybrid webs spun by *Nephilengys* and *Nephilingis* evolved once and are homologous (Fig. 4a). Below we address the three broad questions posed in the Introduction.

**Figure 4.**
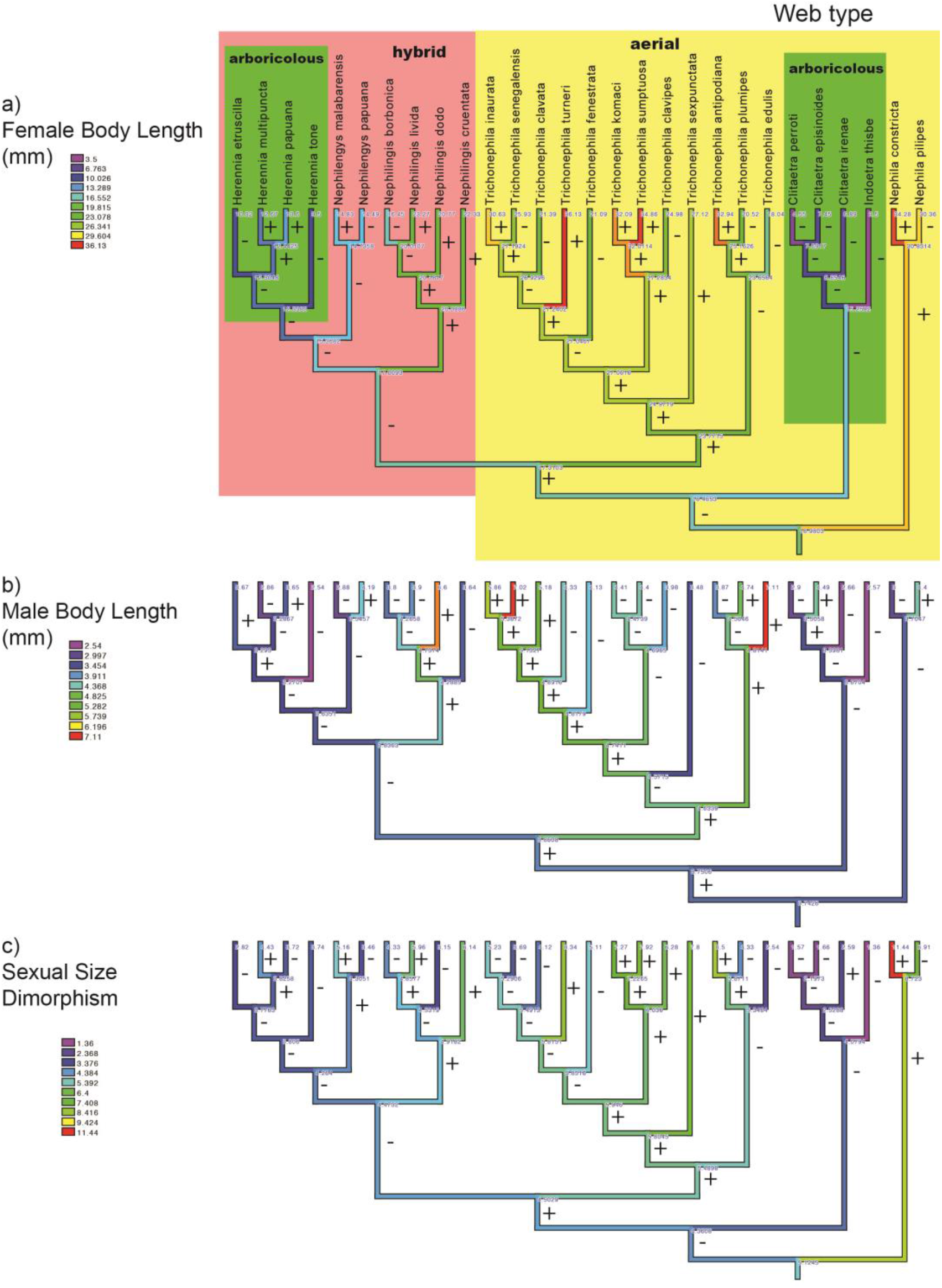
The evolution of body size and sexual size dimorphism (SSD) in nephilid spiders. (a) Female body length shows 26 increases (+) and 28 decreases (-); (b) Male body length has 24 increases and 30 decreases; (c) SSD shows 24 increases and 30 decreases. An ultrametric tree was pruned to contain the 28 taxa with known size variation in both sexes. Sizes and SSD (ratio female/male body length) were optimized on the tree using square change parsimony in Mesquite. Clades are color coded by web type in (a).

1. ***Do male and female size evolve independently, do they evolve towards optima, and how does their evolution affect SSD?. —*** PIC analyses suggest that male and female sizes are independent, both when considering total body length (*r^2^* = 0.0008, 2-tailed *P* = 0.89), as well as carapace length (*r^2^* = 0.016, 2-tailed *P* = 0.52). Brownian motion is the best fit model for nephilid female size and SSD evolution, and OU is the best fit model for male size (Table 2). This suggests that male size evolution, but not female, is driven towards an optimal body size. Squared change parsimony optimizations reveal complex patterns of size evolution in nephilids with numerous increases and decreases (Fig. 4a-b). All terminals and deeper phylogenetic nodes are eSSD except three island *Clitaetra* and *Indoetra* species that independently evolved moderate ratios on Madagascar, Comoros, and Sri Lanka (Fig. 4c). The inferred nephilid root eSSD is 5.1, which is maintained or increased in *Nephila*, and notably in the tropical *Trichonephila, Nephilingis cruentata* (Africa)*, Nephilingis livida* (Madagascar) and *Nephilengys malabarensis* (SE Asia). Conversely, the Australian *T*. *edulis* and *T. plumipes* occupy more temperate areas, and independently evolved smaller eSSD.
2. ***Do nephilids obey Cope’s Rule and the converse Rensch’s Rule?.—*** Ancestral size reconstructions by cladogenetic events (Fig. S5) reveal that female size does not significantly increase (although it trends upward), while male size significantly increases and SSD stagnates, and a linear regression model analysis confirms this (Appendix S1; Fig. S5). These patterns reject Cope’s Rule. Model II regression analysis on phylogenetically independent contrasts data detect no relation between male and female size (Appendix S1, Ma slope = 0.287 (−0.307 to 1.153), *P* = 0.170; see Discussion). Male and female size show no negative allometry, as would be predicted by converse Rensch’s Rule.
3. ***What is the relationship among spider body size and web size, web types, web architecture, and SSD?.—*** Overall, the MCMCglmm analyses reveal that species with larger females spin larger webs (Appendix S1). Female body size significantly differs between web types: species with the largest females spin aerial webs, intermediate sized females spin hybrid webs, and the smallest females spin arboricolous webs (Appendix S1). However, contrary to expectation, MCMCglmm analyses reveal that hybrid webs occupy a larger space than aerial webs (Appendix S1). Squared change parsimony reconstruction of web area (Fig. 5) also shows that *Nephilingis* hybrid webs are largest, that *Nephila* and *Trichonephila* aerial webs are smaller, and that arboricolous webs are smallest, particularly in *Clitaetra*. MCMCglmm analyses suggest that female body size does not affect the ladder index (Appendix S1). Instead, LI in arboricolous webs is significantly higher than in hybrid or aerial webs (Appendix S1); thus, web type affects its “ladderness”. Finally, hub displacement and spider body size or web type are unrelated (Appendix S1). Comparative results furthermore suggest that SSD in arboricolous species is significantly lower than SSD in aerial species, and that SSD in hybrid species does not significantly differ from others (Appendix S1). The convergent evolution of arboricolous webs seems to correspond with a decline in SSD.

**Table 2.** Evolutionary model fitting for nephilid body size and sexual size dimorphism. BM = Brownian motion model; OU = single-optimum Ornstein–Uhlenbeck model; trend = Brownian motion model with a directional trend; FBL = female body length; MBL = male body length; SDI = sexual dimorphism index. P values are from the likelihood ratio test as compared with BM.

| Model | BM | OU | trend |
| --- | --- | --- | --- |
| FBL |  |  |  |
| lnL | -91.672 | -91.672 | -91.370 |
| AIC | 187.344 | 189.344 | 188.740 |
| P value |  | 1.000 | 0.437 |
| MBL |  |  |  |
| lnL | -46.935 | -44.474 | -45.394 |
| AIC | 97.869 | 94.948 | 96.789 |
| P value |  | **0.027** | 0.079 |
| SDI |  |  |  |
| lnL | -61.291 | -60.976 | -61.028 |
| AIC | 126.581 | 127.953 | 128.055 |
| P value |  | 0.428 | 0.468 |

**Figure 5.**
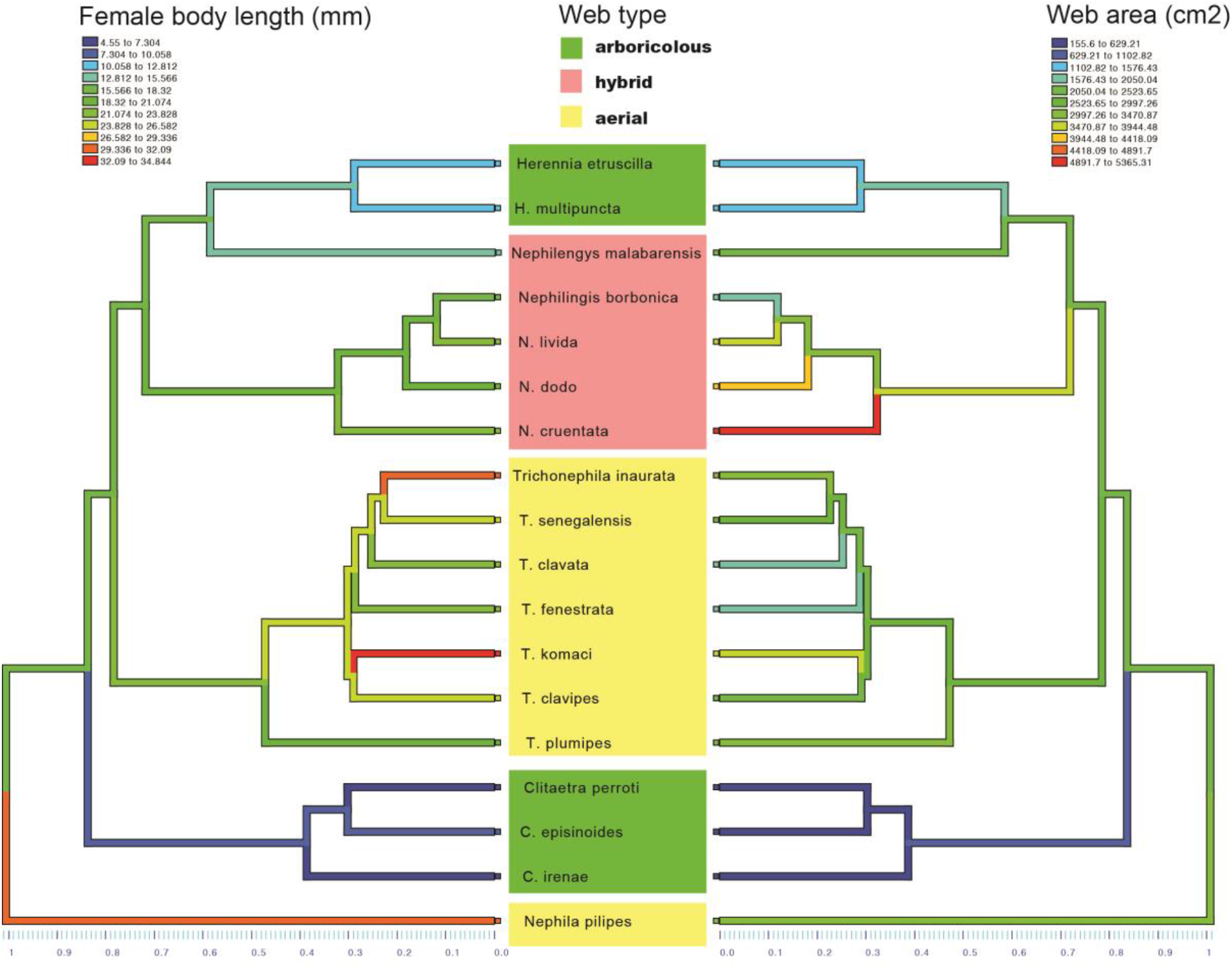
The evolution of female body size and web area. Web types per terminal species are colored as in Figure 4.

## Discussion

We provide a robust new phylogeny of Nephilidae, a model arachnid group for genomic, ecological, biomaterial, and evolutionary research that enables numerous new evolutionary tests to establish patterns in size, web type, and web feature evolution. This phylogenetic foundation also offers an objective rationale for family and genus-level taxonomy.

### Evolutionary Implications

Male and female body sizes evolve independently in Nephilidae, apparently a rare case in animals, and a result that confirms some of the differential equilibrium model predictions of body size evolution (Blanckenhorn 2005). The differential equilibrium model recognizes opposing selection pressures on males and females that, when summed, could push the size of each gender in different directions. Additionally, nephilid gender body sizes evolve independently from one another, but female size is non-directional (Brownian motion), whereas male size is driven towards an optimum (Ornstein-Uhlenbeck model). As suggested by the trendline in Fig. S5, this optimal male body size may lie between 3.2 and 4.9 mm.

As one would expect, female body and web size are strongly and positively correlated. However, even though the largest females spin aerial orb webs, the largest webs— of the hybrid type—are spun by relatively smaller females. The smallest females spin arboricolous, ladder webs. Evolution from aerial towards hybrid webs, as well as the convergent origin of arboricolous webs (once from aerial and once from hybrid webs) have paralleled declines in female body size. Note that since adult males do not spin webs, male web size was not investigated.

Our analyses show that SSD in arboricolous species is smaller than in aerial species, while SSD in hybrid species is not significantly different from either the aerial or the arboricolous clades. In both cases where arboricolous webs evolve, the web site or architecture may constrain gender sizes so that SSD is less. On the other hand, aerial and hybrid webs may facilitate large female body sizes, as they provide less space constraint (Harmer and Herberstein 2009), or enable the capture of larger prey, or easier prey manipulation, or perhaps escape from common substrate dwelling predators. Whatever the reason, the evolutionary result is greater SSD in species on aerial and hybrid webs compared with those on arboricolous webs. Finally, female body size does not affect the web ladder index. Instead, it is the ecology of an arboricolous web that requires a higher ladder index than in hybrid or aerial webs.

### Cope’s and Rensch’s Rules

Several approaches to detect phyletic trends in size yield diverse results. Assessing phyletic size trends in extant and fossil species (Moen 2006) is a strong test. Conventional statistics across ancestor-descendant pairs (Solow and Wang 2008) or model fitting (Monroe and Bokma 2010) are others. Phyletic size increase is supported in the fossil record, but not in extant mammals (Alroy 1998; Monroe and Bokma 2010). Cope’s Rule could be a long-term accumulation of responses to short-term ecological variables. For example, extinct ostracods grew larger as the climate cooled (Hunt and Roy 2006).

Unlike the temperature-related ostracod example, nephilid male phyletic size increase is likely due to gendered fitness advantages of large size at the individual level (Kingsolver and Pfennig 2004). A growing collection of literature has attempted, but so far failed, to converge on general proximate causes, likely due to the fact that male fitness benefits are governed by a mixture of natural and sexual selection (Kuntner and Elgar 2014). Our finding that female body size increase is not significant can likely be attributed to the >100 million year evolutionary timespan. Female gigantism in nephilids is quite obviously an ancient trait (Fig. 6).

**Figure 6.**
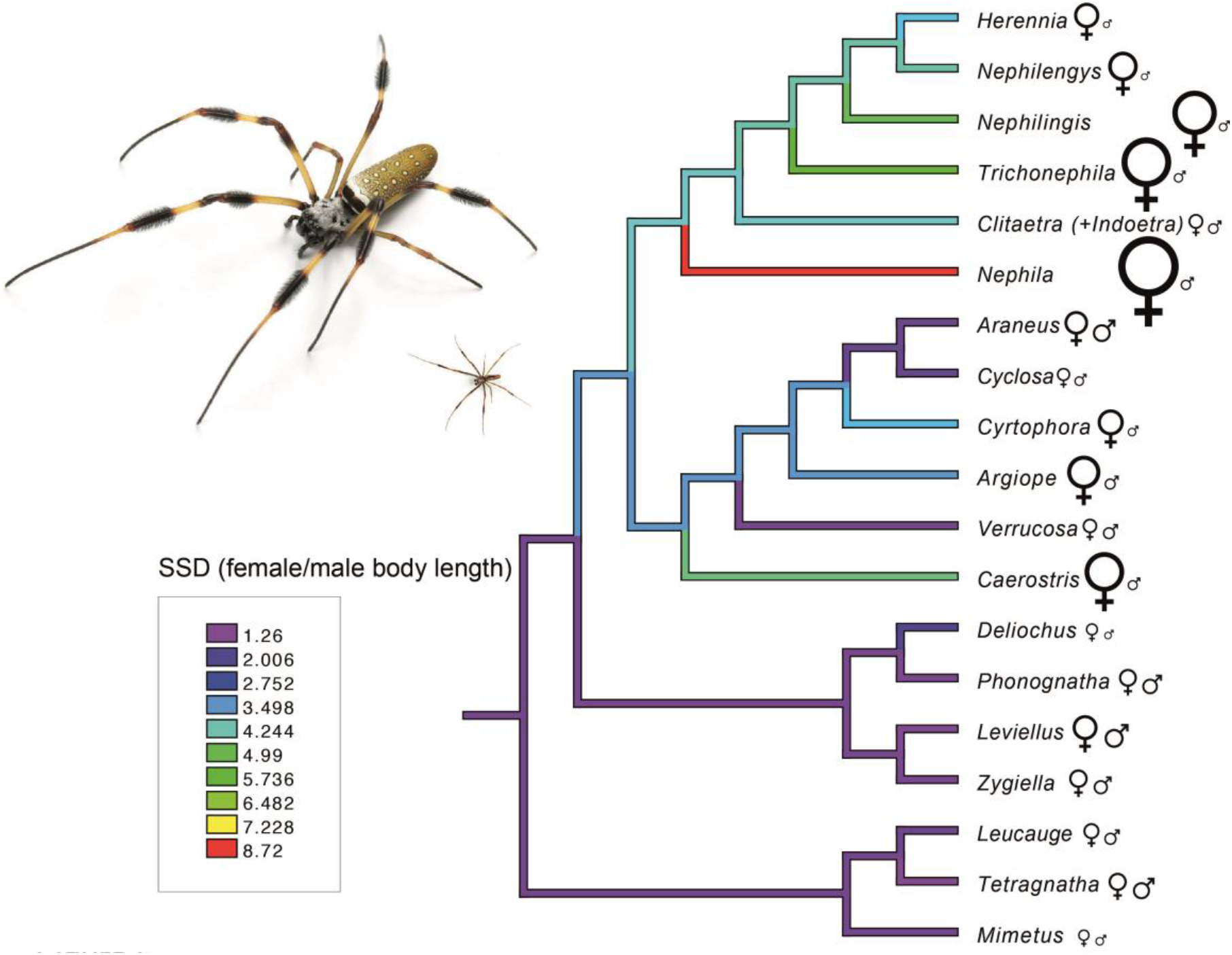
The evolution of sexual size dimorphism (SSD) on a simplified phylogeny. The genus level topology is from AHE analyses, the SSD ratios are optimizations for each nephilid genus (Fig. 4c) or exemplar SSD data for outgroups (Table S3). The reconstruction is linear parsimony in Mesquite. Inset image shows *Trichonephila clavipes* female (left) and male (right). The sizes of female and male symbols correspond to relative total body lengths (Table S3).

Rensch’s Rule predicts that SSD increases with body size in male-biased, and decreases in female-biased SSD species (Abouheif & Fairbairn 1997; Blanckenhorn *et al*. 2007). If female body size changes contribute more to SSD, common in female-biased SSD, then plotting male versus female size, using independent contrasts, should be hypoallometric (Abouheif and Fairbairn 1997; Fairbairn 2005; Foellmer and Moya-Laraño 2007). This pattern, termed converse Rensch’s Rule, should occur in spiders. However, although Abouheif and Fairbairn (1997) corroborated Rensch’s Rule in most male-biased SSD animal taxa, it largely failed in invertebrates (Blanckenhorn *et al*. 2007). Organisms with female biased SSD rarely follow the converse Rensch’s Rule (Webb and Freckleton 2007). In spiders, Foellmer and Moya-Laraño (2007) rejected the converse Rensch’s Rule at interspecific levels, except when analyzing raw (phylogenetically uncorrected) species data. Cheng and Kuntner (2014) rejected the converse Rensch’s Rule after analyzing both raw and phylogenetically corrected data. Nephilid uncorrected data supported the converse Rensch’s Rule, but not phylogenetically corrected data (Kuntner and Cheng 2016). Our results, herein, corroborate that neither Rensch’s Rule, nor its converse, significantly explain sexual size dimorphism in spiders.

### Taxonomy

Our phylogeny has aided the resolution of past taxonomic and classification controversies by recognizing seven genera (*Clitaetra, Indoetra, Herennia, Nephila, Nephilingis, Nephilengys, Trichonephila*) within the family Nephilidae (Table 1). Wheeler et al. (2016) and Dimitrov et al. (2017) treated Nephilinae as a subfamily of Araneidae with no explanation and based on poorly-resolved phylogenies using a limited sampling of genes generally considered to be inappropriate for deep evolutionary inference. We argue that phylogenetic topological results, as well as lineage age, suggest, objectively, that Araneidae, Nephilidae, and Phonognathidae are comparably composed and phylogenetically positioned with respect to other spider families, and should be maintained as independent entities.

Phylogenomic, time-calibrated trees for spider families suggest a range of 37–92 Ma (Garrison et al. 2016). If Nephilidae is a subfamily of Araneidae, our analysis suggests a family age ca. 200 Ma. If such a criterion is applied more broadly across spiders, it would require lumping of diverse spider groups into singular families. Garrison et al.’s chronogram (Fig. 4) at 100 Ma depth would unite not only *Nephila* (Nephilidae) and Araneidae, but also roughly 26 other taxonomically-distinct spider families would disappear. Their exclusion from arachnological taxonomic use is clearly a poor option given their monophyly, age, and evolutionary complexity.

Taxonomic names serve the important purpose of increasing the information content of a classification (Hennig 1965). If Nephilidae, Phonognathidae, and Araneidae are monophyletic lineages, they certainly deserve names. Treating them as subfamilies within an aberrantly ancient (and huge) family does not accomplish this goal and ignores current arachnological practice. First, relatively few of the 4,000+ spider genera are assigned to subfamilies. Subfamilies have been used in particular families, but is generally a rare designation across spiders, and no formal list of subfamilies exists. Arachnologists communicate using family and genus names, and therefore family names increase, in a very practical way, the information content of spider classification. Second, spiders (46,900+ species) are one of the only completely cataloged megadiverse taxonomic groups, consolidated within one global database, the World Spider Catalog (WSC 2017), on which arachnologists universally depend. The WSC uses only two ranks above the species: family and genus, each listed alphabetically. If nephilids and phonognathids are treated as subfamilies within Araneidae, their genera will be sprinkled alphabetically throughout the 173 genera of Araneidae, with no indication of their monophyly or evolutionary distinctiveness. Subfamily definitions and their contents will remain buried in the primary literature. Although the latter reason is only pragmatic, treating Nephilidae and Phonognathidae as families is consistent with arachnological and WSC practice and makes it easier for workers to track future taxonomic changes. In sum, comparable family ages, monophyly, exclusivity, information content, morphological diagnosability, and prevailing community practices all support family rank for Nephilidae and Phonognathidae. Moreover, since phylogenomic analyses all recover Araneidae, Nephilidae, and Phonognathidae as monophyletic, proposing the rank-less name Orbipurae for this clade begins to restore some measure of cladistic hierarchy within the vast and diverse superfamily Araneoidea, and should therefore stimulate comparative work.

### The Placement of *Geratonephila*, and the Origin of Nephilidae

The fossil lineage *Geratonephila* has two possible phylogenetic placements. After evaluating a conservative fossil calibration as a stem nephilid, we find the outcomes of the dating analysis to be realistic and consistent with previous knowledge. Alternatively, a less conservative placement would place *Geratonephila* as the lineage leading to extant *Clitaetra* species from Africa, Madagascar, and Comoros. Interestingly, the size of the described male *Geratonephila burmanica* (3.1 mm) agrees with the phylogenetic reconstruction of the male ancestor of *Clitaetra* (3.4 mm).

Burmese amber is estimated to be 97 to 110 million years old (Poinar and Buckley 2012), corresponding to our hard minimum age boundary. Cruickshank and Ko (2003) attribute the deformation of the Hukawng Basin, the locality of this amber, to the collision of India with Asia, and its subduction beneath the Burma plate. They note that while the amber from these layers is certainly Cretaceous, a more precise age is yet to be determined, which led us to establish a soft boundary at 146 Ma, the beginning of the Cretaceous. More recent estimates suggest this amber age to be at least 100 Ma and of Gondwanan origin (Poinar 2018). Burmese amber has also been found to contain fossil evidence for a more ancient origin of bees (Poinar and Danforth 2006). The origin of Nephilidae may thus be in Cretaceous (contra Kuntner et al. 2013a), and may be Gondwanan after all (Kuntner 2006).

Poinar and Buckley (2012) speculated that *Geratonephila* was a social spider, based only on the co-occurrence of a male and juvenile spider in the same amber inclusion. Insofar as no extant or fossil nephilid shows social behavior, it seems conservative to reject communal behavior in *Geratonephila* (see also Penney 2014). In *Clitaetra*, the first postmolt instar juveniles reside in the webs of their mothers, as do a number of other spider lineages (likely an ancestral state). And perhaps most importantly, male nephilids are known to inhabit female webs for long periods of time as they mate-guard females, waiting for them to mature and become reproductively viable (Kuntner et al. 2009). Together, these facts explain the male and juvenile *Geratonephila burmanica* in the same amber inclusion, as well as our decision to use *Geratonephila* as a stem nephilid for dating.

### Conclusions

Biological rules concerning body sizes do not appear to apply to nephilid spiders. Cope’s Rule predicting phyletic size increase in both sexes, is at best naïve and refuted in this case. SSD complicates the interpretation of Cope’s Rule in lineages where gender sizes evolve independently because the rule applies to both sexes, not just one. Although SSD in nephilids has been proposed to be due primarily to female gigantism, female nephilid size does not significantly increase at macroevolutionary time scales. Even sexually dimorphic gigantism (Kuntner and Elgar 2014), in which both sexes increase in size, but females increase faster, is refuted by our results that only corroborate overall male size increase. The new allometric analyses of log body size also refute Rensch’s Rule, as previously suggested (Kuntner and Cheng 2016).

The emerging picture of the interplay between nephilid female and male body size, extreme sexual size dimorphism (eSSD), as well as the interactions of these variables with web architecture and features, has become more complex. This complexity may be due to the considerable increase in taxon sampling, data density, and phylogenetic accuracy. Patterns that twenty years ago seemed clear based on sparse taxonomic sampling and even sparser quantitative data, now seem much more clade and biology-specific, frustrating both new and old efforts to generalize.

Despite the sometimes conflicting trends seen within Nephilidae, the clade stands as the most extreme example of female-biased sexual size dimorphism among terrestrial animals, as far as we know. Over the years, a large amount of nephilid data has been accumulated on associated phenomena such as fecundity or reduced predation pressure selection for larger females, male-male competition for larger males, or mortality selection for smaller males due to mate searching and avoiding cannibalism, selection for sperm or scramble competition, sexual conflict, genital mutilation or emasculation, and gravity (reviewed in Kuntner and Elgar 2014). However, none of these data can be reliably analyzed in a comparative framework without a stable phylogeny. Comparative perspectives are most powerful when combined with direct experimental data, but here, too, experimental design often depends on phylogenies. Our phylogeny bridges this gap for future comparative studies on this clade of spiders.

Finally, the advent of robust, dated phylogenies can help resolve differing opinions about taxonomic rank by allowing Hennig’s (1965) criterion of rank to be tied to lineage age (“if the absolute rank of categories was linked to their time of origin, just as in geology the sequence of strata in different continents is made comparable by its correlation with specific periods of the earth’s history” p. 115). It seems that spider families, as currently defined, very roughly, tend to be around 40–90 million years old. Though it is unlikely that taxonomic rank will ever be free of subjective opinion, rough norms can be established to guide the application of ranks to evolutionarily distinct monophyletic groups.

## Acknowledgments

We thank Michelle Kortyna, Alyssa Bigelow, Kirby Birch, and Sean Holland for assisting with molecular data collection and bioinformatics analysis, Charles Haddad, and Daiqin Li for field assistance, as well as Simona Kralj-Fišer, Klemen Čandek, Shakira Quiñones-Lebron, Eva Turk, Jutta Schneider, Fritz Vollrath, and George Poinar for helpful discussions.

## Literature Cited

Aberer A.J., Krompass D., Stamatakis A. 2013. Pruning rogue taxa improves phylogenetic accuracy: An efficient algorithm and webservice. Syst. Biol. 62:162–166.

Abouheif E., Fairbairn D.J. 1997. A comparative analysis of allometry for sexual size dimorphism: assessing Rensch’s Rule. Am. Nat. 149:540–562.

Agnarsson I., Coddington J.A., Kuntner M. 2013. Systematics: progress in the study of spider diversity and evolution. In: Penney D., editor. Spider research in the 21st century: Trends and perspectives. Manchester: Siri Scientific Press. p. 58–111.

Alroy J. 1998. Cope’s Rule and the dynamics of body mass evolution in North American fossil mammals. Science 280:731–734.

Babb P.L., Lahens N.F., Correa-Garhwal S.M., Nicholson D.N., Kim E.J., Hogenesch J.B., Kuntner M., Higgins L., Hayashi C.Y., Agnarsson I., Voight B.F. 2017. The *Nephila clavipes* genome highlights the diversity of spider silk genes and their complex expression. Nat. Genet. 49:895–903.

Blackledge T.A., Kuntner M., Agnarsson I. 2011. The form and function of spider orb webs: Evolution from silk to ecosystems. Adv. In Insect Phys. 41:175–262.

Blanckenhorn W.U. 2005. Behavioral causes and consequences of sexual size dimorphism. Ethology 111:977–1016.

Blanckenhorn W.U., Dixon A.F.G., Fairbairn D.J., Foellmer M.W., Gibert P., Linde K. van der, Meier R., Nylin S., Pitnick S., Schoff C., Signorelli M., Teder T., Wiklund C. 2007a. Proximate causes of Rensch’s Rule: Does sexual size dimorphism in arthropods result from sex differences in development time? Am. Nat. 169:245–257.

Blanckenhorn W.U., Meier R., Teder T. 2007b. Rensch’s rule in insects: patterns among and within species. Sex, size and gender roles. Oxford University Press. p. 60–70.

Bokma F., Godinot M., Maridet O., Ladevèze S., Costeur L., SolÉ F., Gheerbrant E., Peigné S., Jacques F., Laurin M. 2016. Testing for Deperet’s Rule (body size increase) in mammals using combined extinct and extant data. Syst. Biol. 65:98–108.

Bond J.E., Garrison N.L., Hamilton C.A., Godwin R.L., Hedin M., Agnarsson I. 2014. Phylogenomics resolves a spider backbone phylogeny and rejects a prevailing paradigm for orb web evolution. Curr. Biol. 24:1765–1771.

Cheng R.-C., Kuntner M. 2014. Phylogeny suggests non-directional and isometric evolution of sexual size dimorphism in argiopine spiders. Evolution 68:1–31.

Cheng R.-C., Kuntner M. 2015. Disentangling the size and shape components of sexual dimorphism. Evol. Biol. 42:223–234.

Coddington J.A., Hormiga G., Scharff N. 1997. Giant female or dwarf male spiders? Nature 385:687–688.

Cruickshank R.D., Ko K. 2003. Geology of an amber locality in the Hukawng Valley, Northern Myanmar. J. Asian Earth Sci. 21:441–455.

Dahl F. 1911. Die Verbreitung der Spinnen spricht gegen eine frühere Landverbindung der Südspitzen unserer Kontinente. Zool. Anz.37:270–282.

Danielson-François A., Hou C., Cole N., Tso I.M. 2012. Scramble competition for moulting females as a driving force for extreme male dwarfism in spiders. Anim. Behav. 84:937–945.

Dimitrov D., Benavides L.R., Arnedo M.A., Giribet G., Griswold C.E., Scharff N., Hormiga G. 2017. Rounding up the usual suspects: a standard target-gene approach for resolving the interfamilial phylogenetic relationships of ecribellate orb-weaving spiders with a new family-rank classification (Araneae, Araneoidea). Cladistics 33:221–250.

Eberhard W.G. 1990. Function and phylogeny of spider webs. Annu. Rev. Ecol. Syst. 21:341–372.

Elgar M.A. 1991. Sexual cannibalism, size dimorphism, and courtship in orb-weaving spiders (Araneidae). Evolution 45:444–448.

Fairbairn D.J. 1997. Allometry for sexual size dimorphism: patterns and process in the coevolution of body size in males and females. Annu. Rev. Ecol. Syst. 28:659–687.

Fairbairn D.J. 2005. Allometry for sexual size dimorphism: Testing two hypotheses for Rensch’s Rule in the water strider *Aquarius remigis*. Am. Nat. 166:S69–S84.

Fairbairn D.J. 2007. Introduction: the enigma of sexual size dimorphism. Sex, size and gender roles. Oxford University Press. p. 1–10.

Fernández R., Hormiga G., Giribet G. 2014. Phylogenomic analysis of spiders reveals nonmonophyly of orb weavers. Curr. Biol. 24:1772–1777.

Fernández R., Kallal R.J., Dimitrov D., Ballesteros J.A., Arnedo M.A., Giribet G., Hormiga G. 2018. Phylogenomics, diversification dynamics, and comparative transcriptomics across the spider tree of life. Curr. Biol. 28:1489–1497.

Foellmer M.W., Moya-Laraño J. 2007. Sexual size dimorphism in spiders: patterns and processes. In: D. J. Fairbairn, W. U. Blanckenhorn and T.S., editor. Sex, size, and gender roles: Evolutionary studies of sexual size dimorphism. Oxford University Press. p. 71–82.

Garrison N.L., Rodriguez J., Agnarsson I., Coddington J.A., Griswold C.E., Hamilton C.A., Hedin M., Kocot K.M., Ledford J.M., Bond J.E. 2016. Spider phylogenomics: untangling the Spider Tree of Life. PeerJ 4:e1719.

Godwin R.L., Opatova V., Garrison N.L., Hamilton C.A., Bond J.E. 2018. Phylogeny of a cosmopolitan family of morphologically conserved trapdoor spiders (Mygalomorphae, Ctenizidae) using Anchored Hybrid Enrichment, with a description of the family, Halonoproctidae Pocock 1901. Mol. Phylogenet. Evol. 126:303–313.

Gould S.J. 1997. Cope’s rule as psychological artefact. Nature 385:199–200.

Gregorič M., Agnarsson I., Blackledge T. A., Kuntner M. 2015. Phylogenetic position and composition of Zygiellinae and *Caerostris*, with new insight into orb-web evolution and gigantism. Zool. J. Linn. Soc. 175:225–243.

Hadfield J.D. 2010. MCMC methods for multi-response generalized linear mixed models: the MCMCglmm R package. J. Stat. Softw. 33:1–22.

Hamilton C.A., Hendrixson B.E., Bond J.E. 2016a. Taxonomic revision of the tarantula genus *Aphonopelma* Pocock, 1901 (Araneae, Mygalomorphae, Theraphosidae) within the United States. Zookeys 2016:1–340.

Hamilton C.A., Lemmon A.R., Lemmon E.M., Bond J.E. 2016b. Expanding anchored hybrid enrichment to resolve both deep and shallow relationships within the spider tree of life. BMC Evol. Biol. 16:212.

Harmer A.M.T., Herberstein M.E. 2009. Taking it to extremes: what drives extreme web elongation in Australian ladder web spiders (Araneidae: *Telaprocera maudae*)? Anim. Behav.78:499–504.

Harmon L.J., Weir J.T., Brock C.D., Glor R.E., Challenger W. 2008. GEIGER: Investigating evolutionary radiations. Bioinformatics 24:129–131.

Head G. 1995. Selection on fecundity and variation in the degree of sexual size dimorphism among spider species (Class Araneae). Evolution 49:776.

Heim N.A., Knope M. 2015. Cope’s rule in the evolution of marine animals. Science 347:867–870.

Hennig W. 1965. Phylogenetic systematics. Annu. Rev. Entomol. 10:97–116.

Higgins L. 2002. Female gigantism in a New Guinea population of the spider *Nephila maculata*. Oikos 99:377–385.

Higgins L., Coddington J., Goodnight C., Kuntner M. 2011. Testing ecological and developmental hypotheses of mean and variation in adult size in nephilid orb-weaving spiders. Evol. Ecol. 25:1289–1306.

Hone D., Benton M. 2005. The evolution of large size: how does Cope’s Rule work? Trends Ecol. Evol. 20:4–6.

Hormiga G., Griswold C.E. 2014. Systematics, phylogeny, and evolution of orb-weaving spiders. Annu. Rev. Entomol. 59:487–512.

Hormiga G., Scharff N., Coddington J.A. 2000. The phylogenetic basis of sexual size dimorphism in orb-weaving spiders (Araneae, Orbiculariae). Syst. Biol. 49:435–62.

Hunt G., Roy K. 2006. Climate change, body size evolution, and Cope’s Rule in deep-sea ostracodes. Proc. Natl. Acad. Sci. 103:1347–52.

Kallal R.J., Fernández R., Giribet G., Hormiga G. 2018. A phylotranscriptomic backbone of the orb-weaving spider family Araneidae (Arachnida, Araneae) supported by multiple methodological approaches. Mol. Phylogenet. Evol. 126:129–140.

Kallal R.J., Hormiga G. 2018. Systematics, phylogeny and biogeography of the Australasian leaf-curling orb-weaving spiders (Araneae: Araneidae: Zygiellinae), with a comparative analysis of retreat evolution. Zool. J. Lin. Soc. XX:1–87.

Katoh K., Standley D.M. 2013. MAFFT multiple sequence alignment software version 7: Improvements in performance and usability. Mol. Biol. Evol. 30:772–780.

Kingsolver J.G., Pfennig D.W. 2004. Individual-level selection as a cause of Cope’s rule of phyletic size increase. Evolution 58:1608–1612.

Kuntner M. 2005. A revision of *Herennia* (Araneae: Nephilidae: Nephilinae), the Australasian “coin spiders”. Invertebr. Syst.19:391–436.

Kuntner M. 2006. Phylogenetic systematics of the Gondwanan nephilid spider lineage Clitaetrinae (Araneae, Nephilidae). Zool. Scr.35:19–62.

Kuntner M. 2007. A monograph of *Nephilengys*, the pantropical ‘hermit spiders’ (Araneae, Nephilidae, Nephilinae). Syst. Entomol. 32:95–135.

Kuntner M. 2017. Nephilidae. In: Ubick D., Paquin P., Cushing P.E., Roth V., editors. Spiders of North America: An identification manual (second edition). p. 191–192.

Kuntner M., Agnarsson I. 2011. Biogeography and diversification of hermit spiders on Indian Ocean islands (Nephilidae: *Nephilengys*). Mol. Phylogenet. Evol. 59:477–488.

Kuntner M., Arnedo M.A., Trontelj P., Lokovšek T., Agnarsson I. 2013. A molecular phylogeny of nephilid spiders: Evolutionary history of a model lineage. Mol. Phylogenet. Evol.69:961–979.

Kuntner M., Cheng R.-C. 2016. Evolutionary pathways maintaining extreme female-biased sexual size dimorphism: convergent spider cases defy common patterns. In: Pontarotti P., editor. Evolutionary biology: Convergent evolution, evolution of complex traits, concepts and methods. Cham: Springer International Publishing. p. 121–133.

Kuntner M., Cheng R.-C., Kralj-Fišer S., Liao C.-P., Schneider J.M., Elgar M.A. 2016. The evolution of genital complexity and mating rates in sexually size dimorphic spiders. BMC Evol. Biol. 16:242.

Kuntner M., Coddington J.A. 2009. Discovery of the largest orbweaving spider species: The evolution of gigantism in *Nephila*. PLoS One 4:2–6.

Kuntner M., Coddington J.A., Hormiga G. 2008. Phylogeny of extant nephilid orb-weaving spiders (Araneae, Nephilidae): testing morphological and ethological homologies. Cladistics 24:147–217.

Kuntner M., Coddington J.A., Schneider J.M. 2009. Intersexual arms race? Genital coevolution in nephilid spiders (Araneae, Nephilidae). Evolution 63:1451–1463.

Kuntner M., Elgar M.A. 2014. Evolution and maintenance of sexual size dimorphism: Aligning phylogenetic and experimental evidence. Front. Ecol. Evol. 2:1–8.

Kuntner M., Gregorič M., Li D. 2010. Mass predicts web asymmetry in *Nephila* spiders. Naturwissenschaften 97:1097–1105.

Kuntner M., Zhang S., Gregorič M., Li D. 2012. *Nephila* female gigantism attained through post-maturity molting. J. Arachnol. 40:345–347.

Legendre P. 2014. lmodel2: Model II Regression. R package version 1.7–2.

Lemmon A.R., Emme S.A., Lemmon E.M. 2012. Anchored hybrid enrichment for massively high-throughput phylogenomics. Syst. Biol. 61:727–744.

Lupše N., Cheng R.-C., Kuntner M. 2016. Coevolution of female and male genital components to avoid genital size mismatches in sexually dimorphic spiders. BMC Evol. Biol.16:161.

Maddison W.P., Evans S.C., Hamilton C.A., Bond J.E., Lemmon A.R., Lemmon E.M. 2017. A genome-wide phylogeny of jumping spiders (Araneae, Salticidae), using anchored hybrid enrichment. Zookeys 2017:89–101.

Maddison W.P., Maddison D.R. 2015. Mesquite: a modular system for evolutionary analysis. Version 3.04. URL http://mesquiteproject.org.

Meyer M., Kircher M. 2010. Illumina sequencing library preparation for highly multiplexed target capture and sequencing. Cold Spring Harb. Protoc. 2010:pdb.prot5448–pdb.prot5448.

Mirarab S., Warnow T. 2015. ASTRAL-II: coalescent-based species tree estimation with many hundreds of taxa and thousands of genes. Bioinformatics 31:i44–i52.

Moen D.S. 2006. Cope’s rule in cryptodiran turtles: Do the body sizes of extant species reflect a trend of phyletic size increase? J. Evol. Biol. 19:1210–1221.

Monroe M.J., Bokma F. 2010. Little evidence for Cope’s rule from Bayesian phylogenetic analysis of extant mammals. J. Evol. Biol. 23:2017–2021.

Moya-Laraño J., Halaj J., Wise D.H. 2002. Climbing to reach females: Romeo should be small. Evolution 56:420–425.

Moya-Laraño J., Vinković D., Allard C.M., Foellmer M.W. 2009. Optimal climbing speed explains the evolution of extreme sexual size dimorphism in spiders. J. Evol. Biol. 22:954–963.

Nguyen L.T., Schmidt H.A., Von Haeseler A., Minh B.Q. 2015. IQ-TREE: A fast and effective stochastic algorithm for estimating maximum-likelihood phylogenies. Mol. Biol. Evol. 32:268–274.

Penney D. 2014. Predatory behaviour of Cretaceous social orb-weaving spiders: comment. Hist. Biol. 26:132–134.

Poinar G. 2018. Burmese amber: evidence of Gondwanan origin and Cretaceous dispersion. Hist. Biol. 2963:1–6.

Poinar G., Buckley R. 2012. Predatory behaviour of the social orb-weaver spider, *Geratonephila burmanica* n. gen., n. sp. (Araneae: Nephilidae) with its wasp prey, *Cascoscelio incassus* n. gen., n. sp. (Hymenoptera: Platygastridae) in early Cretaceous Burmese amber. Hist. Biol. 24:519–525.

Poinar G.O., Danforth B.N. 2006. A fossil bee from early cretaceous burmese amber. Science 314:614–614.

Prum R.O., Berv J.S., Dornburg A., Field D.J., Townsend J.P., Lemmon E.M., Lemmon A.R. 2015. A comprehensive phylogeny of birds (Aves) using targeted next-generation DNA sequencing. Nature 526:569–573.

Ramos M., Coddington J.A., Christenson T.E., Irschick D.J. 2005. Have male and female genitalia coevolved? A phylogenetic analysis of genitalic morphology and sexual size dimorphism in web-building spiders (Araneae: Araneoidea). Evolution 59:1989–1999.

Rensch B. 1948. Histological changes correlated with evolutionary changes of body size. Evolution 2:218–230.

Rokyta D.R., Lemmon A.R., Margres M.J., Aronow K. 2012. The venom-gland transcriptome of the eastern diamondback rattlesnake (*Crotalus adamanteus*). BMC Genomics13:312.

Scharff N., Coddington J.A. 1997. A phylogenetic analysis of the orb-weaving spider family Araneidae (Arachnida, Araneae). Zool. J. Linn. Soc. 120:355–434.

Schneider J.M., Elgar M.A. 2001. Sexual cannibalism and sperm competition in the golden orb-web spider *Nephila plumipes* (Araneoidea): female and male perspectives. Behav. Ecol.12:547–552.

Solow A.R., Wang S.C. 2008. Some problems with assessing Cope’s Rule. Evolution 62:2092–2096.

Stanley S. 1973. An explanation for Cope’s Rule. Evolution 27:1–26.

Starrett J., Derkarabetian S., Hedin M., Bryson R.W., McCormack J.E., Faircloth B.C. 2017. High phylogenetic utility of an ultraconserved element probe set designed for Arachnida. Mol. Ecol. Resour. 17:812–823.

Tamura K., Battistuzzi F.U., Billing-Ross P., Murillo O., Filipski A., Kumar S. 2012. Estimating divergence times in large molecular phylogenies. Proc. Natl. Acad. Sci. 109:19333–19338.

Vidergar N., Toplak N., Kuntner M. 2014. Streamlining DNA barcoding protocols: Automated DNA extraction and a new *cox1* primer in arachnid systematics. PLoS One 9:e113030.

Vollrath F., Parker G.A. 1992. Sexual dimorphism and distorted sex ratios in spiders. Nature 360:156–159.

Waller J.T., Svensson E.I. 2017. Body size evolution in an old insect order: No evidence for Cope’s Rule in spite of fitness benefits of large size. Evolution 71:2178–2193.

Webb T.J., Freckleton R.P. 2007. Only half right: species with female-biased sexual size dimorphism consistently break Rensch’s Rule. PLoS One 2:e897.

Wheeler W.C., Coddington J.A., Crowley L.M., Dimitrov D., Goloboff P.A., Griswold C.E., Hormiga G., Prendini L., Ramírez M.J., Sierwald P., Almeida-Silva L., Alvarez-Padilla F., Arnedo M.A., Benavides Silva L.R., Benjamin S.P., Bond J.E., Grismado C.J., Hasan E., Hedin M., Izquierdo M.A., Labarque F.M., Ledford J., Lopardo L., Maddison W.P., Miller J.A., Piacentini L.N., Platnick N.I., Polotow D., Silva-Dávila D., Scharff N., Szűts T., Ubick D., Vink C.J., Wood H.M., Zhang J. 2017. The spider tree of life: phylogeny of Araneae based on target-gene analyses from an extensive taxon sampling. Cladistics 33:574–616.

WSC. 2017. World Spider Catalog version 18.0. Available from http://wsc.nmbe.ch.

Wunderlich J. 1986. Spinnenfauna Gestern und Heute. Wiesbaden: Bauer Verlag.

Wunderlich J. 2004. The fossil spiders (Araneae) of the families Tetragnathidae and Zygiellidae n. stat. in Baltic and Dominican amber, with notes on higher extant and fossil taxa. Beitr. Araneol. 3:899–955.

